# Enhancing Embryo Stage Classification with Multi-Focal Plane Imaging and Deep Learning

**DOI:** 10.1101/2025.03.21.644547

**Authors:** Aissa Benfettoume Souda, Wistan Marchadour, Souaad Hamza-Cherif, Jean-Marie Guyader, Marwa Elbouz, Fréderic Morel, Aurore Perrin, Mohammed El Amine Bechar, Nesma Settouti

## Abstract

**Purpose:** Accurately identifying embryo developmental stages is crucial for improving success rates in in vitro fertilization (IVF). Traditional embryo assessment relies on 2D imaging with a single focal plane, which may overlook critical morphological details and lead to misclassification. This study investigates whether incorporating depth information through multi-focal plane imaging can enhance the accuracy of embryo stage classification.

**Methods:** We compared 2D and 3D convolutional neural network (CNN) architectures trained on a time-lapse embryo dataset containing seven focal planes. The models were evaluated based on their classification performance across different embryo developmental stages. Additionally, we used Gradient-weighted Class Activation Mapping (Grad-CAM) to analyze model attention and interpretability.

**Results:** The findings demonstrate that 3D CNN models leveraging multi-focal plane data significantly outperform their 2D counterparts, particularly in complex stages such as Morula formation (tM) and expanded blastocyst (tSB, tB, and tEB). Grad-CAM visualization confirms that the models focus on relevant morphological structures, further validating the advantages of volumetric modeling.

**Conclusion:** Multi-focal plane imaging enhances embryo stage classification accuracy by providing richer morphological information. This study underscores the potential of volumetric modeling in embryo assessment and paves the way for extending this approach to full 3D time-lapse analysis.

## 1 Introduction

In vitro fertilization (IVF) is a cornerstone of assisted reproductive technologies (ART), involving fertilization of an oocyte by sperm outside the body, followed by embryo transfer to the uterus to achieve pregnancy. IVF offers hope to couples facing infertility and to individuals experiencing challenges conceiving naturally [1]. In the U.S., approximately 389,993 IVF cycles were performed in 2022, leading to more than 91,771 live births. However, the average success rate per cycle remains limited, with only about 37.3% leading to a live birth [2]. This moderate success rate, combined with the significant emotional, physical, and financial burden of IVF, underscores the importance of improving embryo selection to maximize pregnancy outcomes.

The success of IVF largely depends on selecting viable embryos, traditionally based on morphological assessments from time-lapse imaging (TLI) or standard microscopy [3]. However, these evaluations are subjective and lack scalability, prompting the integration of AI to enhance precision and reproducibility in embryo quality assessment. By leveraging the rich spatial and temporal data from TLI and static images, AI-driven approaches stratify embryonic quality and predict developmental potential, implantation success, and clinical pregnancy outcomes. Despite significant progress, challenges remain due to variations in AI methodologies, dataset limitations, and the need for standardized evaluation criteria. Addressing these challenges is key to improving automated embryo selection and clinical outcomes.

Recent studies have explored the use of Deep Learning (DL) in embryo evaluation, achieving promising results in predicting developmental stages and pregnancy outcomes [4–6]. However, with early efforts centering on static, single-plane images to predict IVF outcomes. For instance, models trained on 2D static images have demonstrated potential in assessing pregnancy likelihood [4] and curating datasets of blastocyst images for training DL classifiers [5]. A recent review highlighted that DL-based models often outperforms traditional assessments, although largely based on 2D historical data [7]. These initiatives have shown promise in simplifying embryo evaluation, yet their reliance on single-plane data limits the capture of deeper structural details, signaling a need for broader exploration.

Building on static methods, TLI has introduced temporal dynamics to embryo assessment, typically relying on a few focal planes. A dataset of TLI sequences was examined for blastocyst prediction, testing single-frame and multi-frame architectures [8], but its multi-frame approach still hinges on limited focal depth, missing broader spatial detail. Another developed a scoring system for cohort evaluation based on TLI [9], yet its reliance on sparse focal planes curbs its structural insight. One analysis scored developmental potential across multiple stages with vast TLI data [10], offering robust temporal coverage, though its focus on sequence over depth limits 3D potential. Meanwhile, a technique enhanced image clarity through multi-focus fusion [11], a promising step, but it prioritizes enhancement over full volumetric modeling. Collectively, these efforts advance automation and temporal analysis, yet their minimal use of focal planes restricts a comprehensive embrace of the embryo’s spatial structure, leaving significant room for deeper exploration.

To address this gap, our research explores the impact of multi-focal depth in embryo assessment. Leveraging existing datasets [12], we propose a 3D CNN architecture based on ResNet to track embryonic development. Unlike previous studies that primarily rely on single-plane (2D) imaging, our approach systematically compares both single-plane and multi-plane (3D) representations to assess how the integration of depth information influences model performance in classifying 15 embryonic developmental stages. Additionally, we incorporate Gradient-weighted Class Activation Mapping (Grad-CAM) to interpret the model’s decision-making process and analyze feature importance across different focal depths. By integrating volumetric data and explainability techniques, we aim to refine automated embryo selection and provide a more comprehensive framework for embryo evaluation, bridging a critical gap in DL-driven reproductive medicine.

### Highlights

- First systematic comparison of single-plane (2D) vs. multi-plane (3D) imaging in embryo assessment.
- Demonstration of the added value of volumetric data to enhance classification accuracy and developmental tracking.
- Integration of Grad-CAM to provide interpretability and highlight critical features used in model predictions.
- Contribution to AI-driven reproductive medicine by improving automation, transparency, and decision-making in embryo selection.

The remainder of this paper is organized as follows: Section 2 details the methodology, including data collection, the focal plane strategy. The proposed deep learning architectures are presented in Section 3, both 2D and 3D ResNet-based models designed to classify embryo developmental stages are presented, alongside the integration of Grad-CAM for model interpretability. Section 4 describes the experimental setup, outlining data preparation, hyperparameter tuning, and training protocols. We then report and analyze the classification results, highlighting the impact of depth information on model performance. Finally, Section 5 concludes the paper with a discussion of the study’s main findings and proposes future directions for enhancing AI-driven embryo evaluation in clinical practice.

## 2 Methodology

In this section, we describe the embryo time-lapse dataset used, which offers detailed stage annotations and multi-focal imaging. We then explain the focal plane acquisition strategy, emphasizing how depth information enhances the morphological representation of developing embryos.

### 2.1 Data collection

We used a publicly available dataset [12] of 704 time-lapse videos of developing embryos, each recorded at seven focal planes, for a total of 2.4 million images. The dataset includes detailed annotations of 16 developmental phases, which are summarized in Table 1. These stages measure the embryo’s progression from fertilization to blastocyst formation, including early events like polar body extrusion and pronuclei appearance, followed by cleavage phases (e.g., 2-cell, 4-cell, 8-cell), compaction, morula formation, and blastocyst milestones such as cavity initiation and expansion Fig. 1. This granular annotation supports the training CNN models to classify stages with high precision.

**Table 1.** Description of stages and Their Counts in the used Dataset.

| Phase | Description | Count in Dataset |
| --- | --- | --- |
| tPB2 | Second polar body extrusion | 8 895 |
| tPNa | Pronuclei appearance | 43 244 |
| tPNf | Pronuclei fading | 6 793 |
| t2 | 2-cell cleavage stage | 29 189 |
| t3 | 3-cell cleavage stage | 5 127 |
| t4 | 4-cell cleavage stage | 29 177 |
| t5 | 5-cell cleavage stage | 8 045 |
| t6 | 6-cell cleavage stage | 8 366 |
| t7 | 7-cell cleavage stage | 10 531 |
| t8 | 8-cell cleavage stage | 32 602 |
| t9+ | 9 or more cell stage (pre-compaction) | 51 112 |
| tM | Morula formation | 17 084 |
| tSB | Start of blastulation (cavity initiation) | 17 244 |
| tB | Blastocyst formation | 10 387 |
| tEB | Expanded blastocyst | 19,535 |
| tHB | Hatching blastocyst | 97 |

**Fig. 1.**
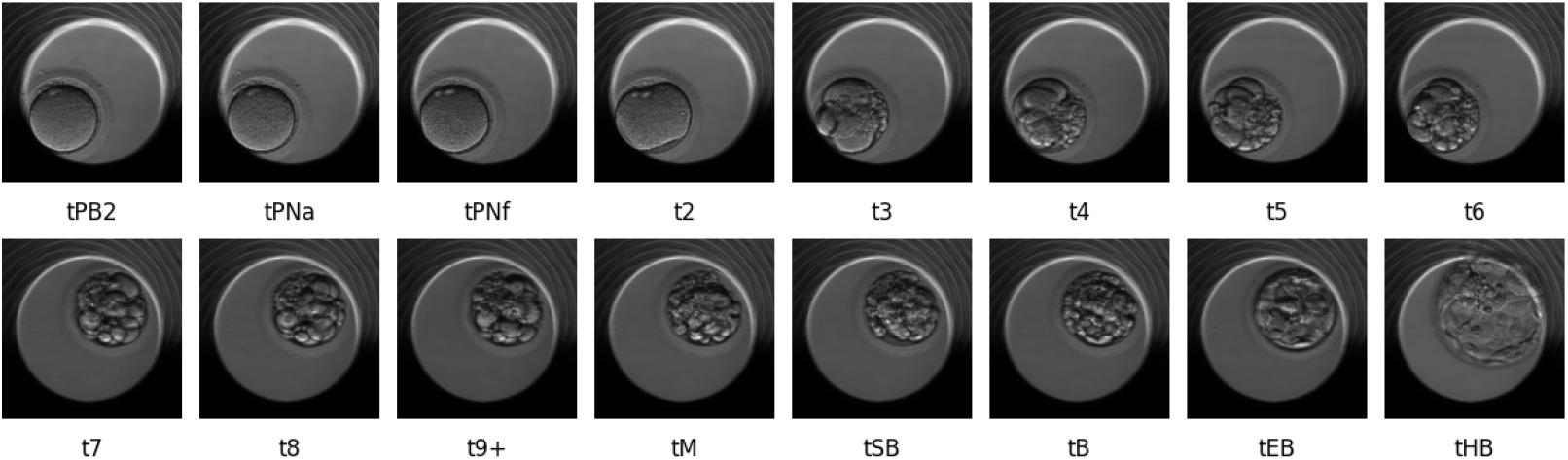
Illustration of the 16 development stages presented in the used dataset.

### 2.2 Focal Plane

Since embryos are three-dimensional structures, imaging at multiple focal planes is crucial for capturing fine morphological details. Each embryo in the dataset was recorded across seven focal planes to provide a more complete representation of its structure Fig. 2. These focal planes are as follows:

**Fig. 2.**
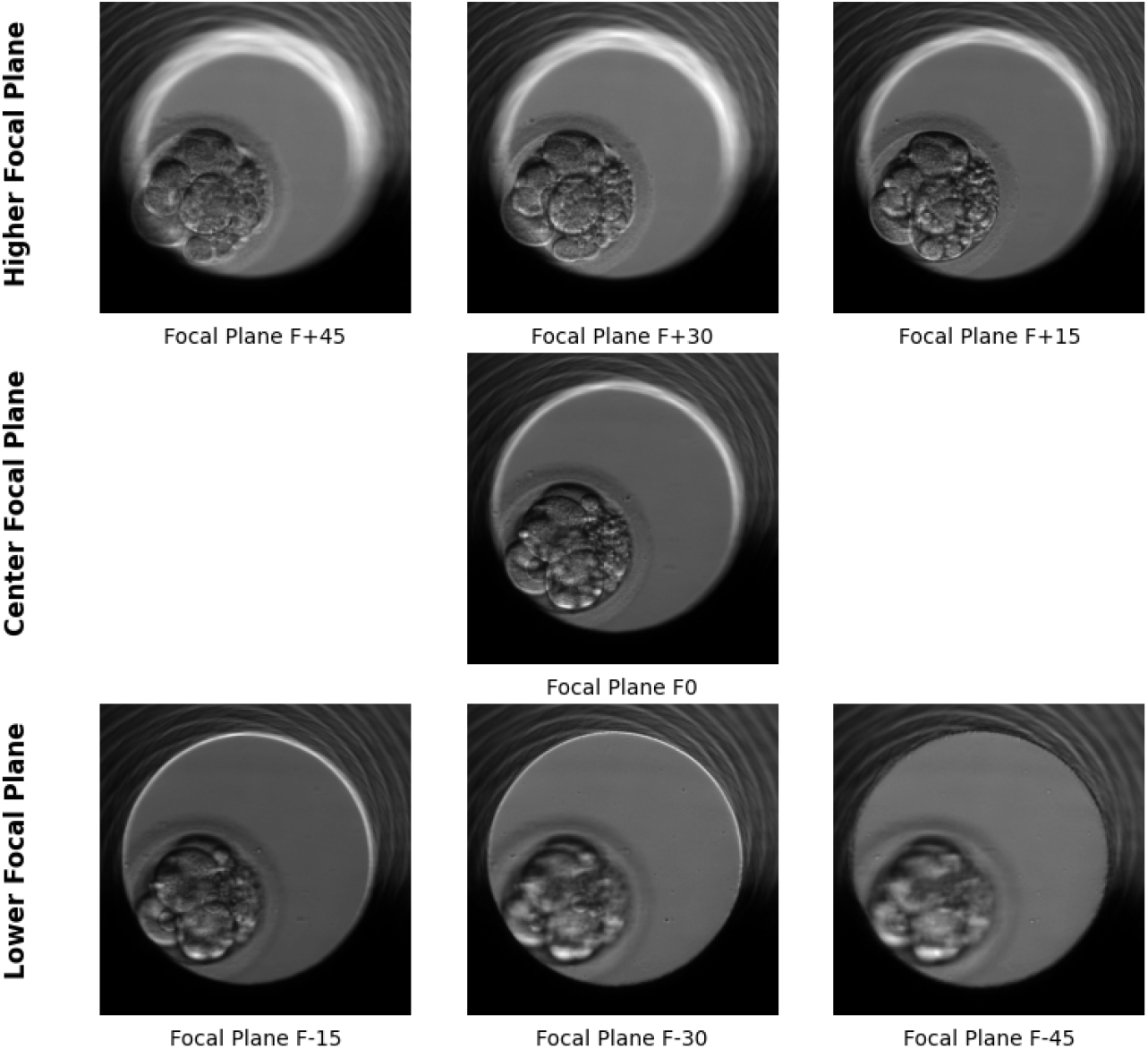
An Illustration Example of the seven focal planes presented in this dataset ranging from F-45 to F+45.

- F-45, F-30 and F-15: are lower focal planes, capturing deeper layers of the embryo.
- F0: is The central focal plane, providing the most balanced and sharpest image.
- F+15, F+30 and F+45: are higher focal planes, capturing the outer layers of the embryo.

By acquiring images at multiple focal depths, the dataset enables more precise identification of key cellular events, reducing the risk of missing important morphological features due to plane selection. Each stage is captured across the seven focal planes, allowing for a detailed 3D reconstruction of morphological changes over time.

## 3 Models design and Explainability

To exploit the dataset’s multi-plane richness, we propose a method employing CNNs to classify embryo developmental stages, comparing 2D single-focal-plane and 3D multi-focal-plane configurations. Unlike prior approaches that rely on sparse focal stacks or mischaracterize 3D as temporal sequences, our method explores the embryo’s true spatial depth, leveraging all seven focal planes (F-45 to F+45) alongside a single-plane baseline (F0) to assess the impact of volumetric data on classification performance Fig. 3.

**Fig. 3.**
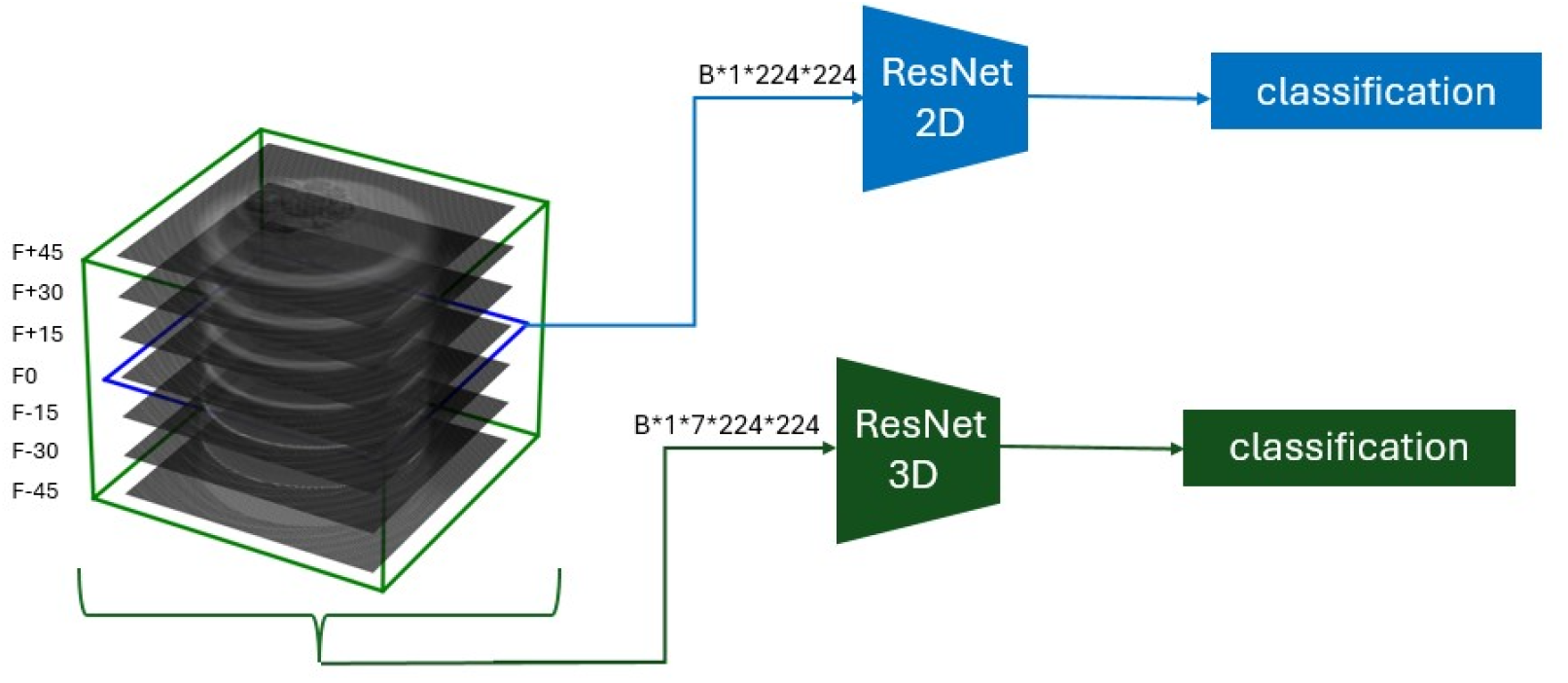
Proposed Method for classifying Embryo stages

### 3.1 Single Focale Plane (2D)

We employ ResNet-18 and ResNet-50 [13], well-known architectures pretrained on ImageNet Dataset [14], to process images from the central focal plane (F0) only. This setup mirrors traditional 2D CNN approaches in embryo analysis, where each time point is treated as an independent grayscale image for developmental stage classification. While computationally efficient and effective in capturing surface-level morphology, this method disregards depth information from the other six planes, potentially limiting its ability to detect subtle 3D structural variations.

### 3.2 Multi Focale Plane (3D)

We extend ResNet-18 and ResNet-50 into 3D CNNs by replacing 2D convolutional kernels with 3D equivalents, enabling volumetric processing.Those architectures were inilazed with random weight. Images from all seven focal planes (F-45, F-30, F-15, F0, F+15, F+30 and F+45) are stacked into a 7×224×224 input tensor, where the depth dimension represents the focal stack. This architecture integrates lower (F-45 to F-15), central (F0), and higher (F15 to F45) focal data, capturing both intra-plane features and inter-plane relationships. By leveraging the full 3D morphology of the embryo, this approach aims to improve classification accuracy and robustness compared to the 2D baseline.

### 3.3 Explainability of CNNs

To better understand the decision process of each model, and estimate the differences between single and multi-focal approaches, we propose to use eXplainable Artificial Intelligence (XAI) techniques. Among the numerous existing XAI concepts [15], we specifically choose to inspect Gradient-weighted Class Activation Mapping (Grad-CAM) [16], which belongs to the saliency-based methods, a category yielding more visual and comprehensible results.

The GradCAM process consists of a weighted combination between the advanced features learned by the network, and gradient backpropagation estimating the contribution of each model parameter to the prediction. It is considered one of the most accurate saliency-based XAI method for model representation, in addition to providing easy-to-read heatmaps. This choice of algorithm is also motivated by the expectation of only one important region per input on this specific application (i.e. the embryo cells), a case for which GradCAM is most efficient. However, this technique is not without a few drawbacks. First, the saliency maps are sensible to a user-specific convolution layer choice, as the only hyper-parameter. Then, because a resizing step is applied from a smaller scale, it causes blurry information on the final maps (as seen below in 4.5), as well as better visual clarity. Therefore, GradCAM induces a quality trade-off, observed in this study.

For the XAI experiment, we focus solely on test images (unused during training) that are correctly classified by all four models, an important factor to avoid faulty contributions, and better understand which elements are deemed important by the networks. From the available subset corresponding to these criteria, we randomly select 5000 images, visually inspect their respective saliency maps, and compare the results across model architectures, classes, and dimensionality. As for the choice of layer on which the computation is performed, we follow the original paper guidelines [16], and select the last convolution step before the final fully-connected layer, for each models.

## 4 Experiments and results

In this section, we present the experiments conducted to evaluate the performance of our models in classifying embryo developmental phases. Using two GPU T4 graphics cards and the PyTorch framework, we preprocess the dataset, train both 2D and 3D models, and assess their performance using key evaluation metrics.

### 4.1 Data preparation

To ensure robust analysis of embryo development, the raw dataset underwent preprocessing to address inconsistencies and enhance model training reliability. Fig. 4 illustrates the original dataset’s distribution across 16 phases, revealing uneven image counts. Specific preprocessing steps include dataset Balancing: Due to the high diversity of the dataset, and due to the limited number of images (only 96) in the last developmental phase (tHB), we excluded this phase and we selected 5,000 images for each of the remaining 15 developmental phases to ensure a balanced representation across all phases. The new balanced dataset consist of 75k images for the 15 development phases was divided into three splits: 56% training, 14% validation, and 30% testing. Table 2 illustrate the dataset split is shown below.

**Table 2.** Dataset repartition.

| Split | Number of Images |
| --- | --- |
| Train | 42000 |
| Validation of Train | 10500 |
| Test | 22500 |

**Fig. 4.**
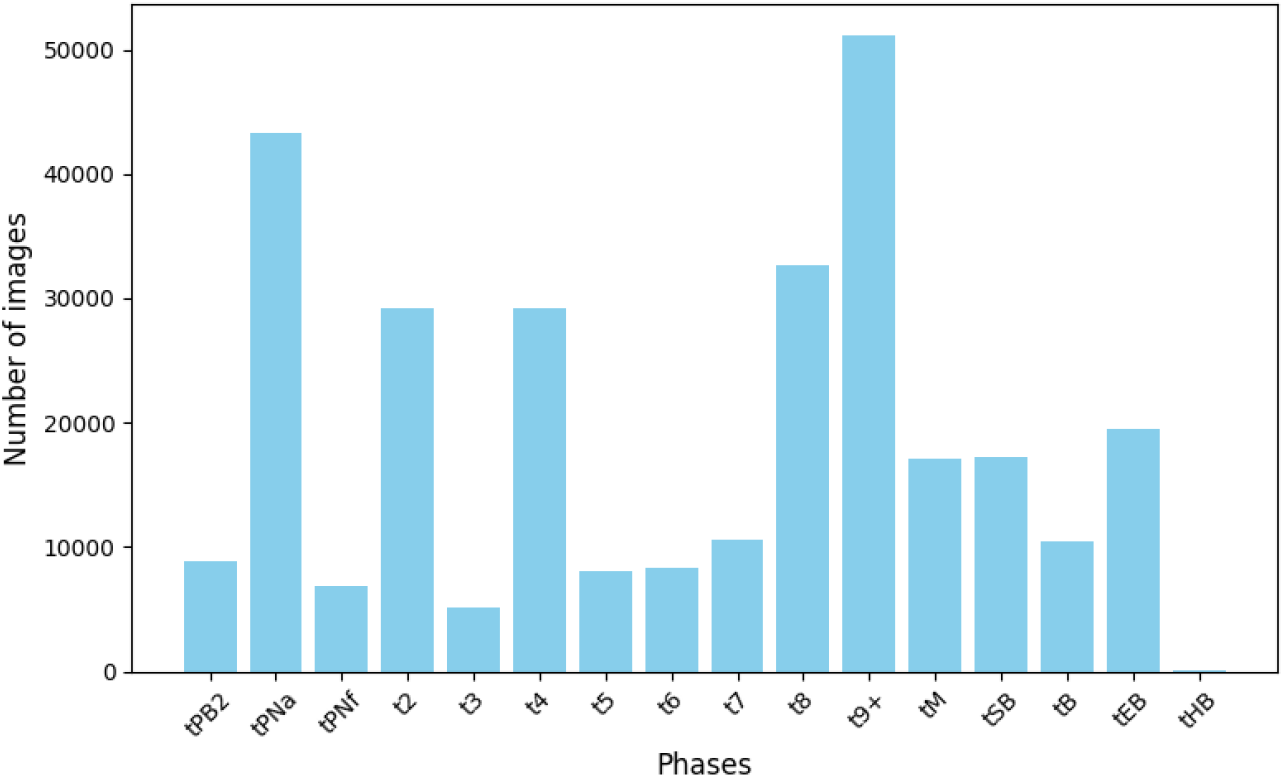
Illustration for distribution of phases across dataset

### 4.2 Hyperparameter settings

To ensure the reliability and consistency of our experiments, we used the same hyper-parameter settings across all models. This approach allows for fair comparisons and reproducible results. The selected hyper-parameters are summarized in Table 3.

**Table 3.** Hyperparameters used for 2D and 3D models.

| Hyperparameter | Description |
| --- | --- |
| Image Size | 224x224 |
| Normalization | *mean, std* |
| Optimizer | Adam |
| Learning Loss | Cross-Entropy Loss |
| Learning rate | $1 \times 10^{-3}$ |
| Batch Size | 16 |
| Epoch | 100 |

### 4.3 Performance metrics

The performance of the 2D and 3D models was compared using metrics such as accuracy, precision, recall, and F1-score. The equations for these metrics are as follows:

**Accuracy**: measures the overall proportion of correct predictions and is calculated as:

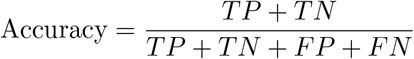

where *TP* is true positives, *TN* is true negatives, *FP* is false positives, and *FN* is false negatives.

**Precision**:Precision measures the proportion of correct positive predictions.

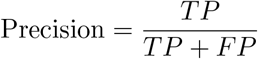

**Recall**:Recall measures the proportion of actual positives that were correctly identified by the model.

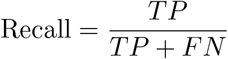

**F1-score**:The F1-score is the harmonic mean of precision and recall, providing a balance between the two metrics.

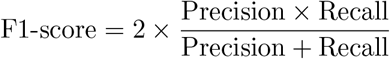

These metrics were used to evaluate and compare the performance of both the 2D and 3D models, helping to assess the models’ effectiveness in classifying the developmental phases of the embryos.

### 4.4 Results

The 2D single-focal-plane (**2D-ResNet18** and **2D-ResNet50**) and 3D multi-focal-plane (**3D-ResNet18** and **3D-ResNet50**) models were evaluated on a test set from the balanced dataset, with performance summarized in Table 4. The **2D-ResNet18** achieved an average F1-score of 0.80, performing well on early stages like tPB2 (0.84 F1-Score) but weaker on t9+ (0.68 F1-Score) due to its single-plane limitation. The **2D-ResNet50** averaged an F1-score of 0.81, enhancing t5 (0.81 F1-Score) yet still struggling with t9+ (0.75 F1-Score), showing modest gains from deeper architecture. In contrast, the 3D models, leveraging all seven focal planes, excelled—**3D-ResNet18** with an average F1-score of 0.87 and **3D-ResNet50** at 0.86—improving complex stages like tEB (F1-Score) and tM (0.84 F1-Score) by capturing depth-wise details. Fig. 5 highlights this 3D advantage across stages.

**Table 4.** Classification Performance of 2D and 3D CNN Models on Test Set.

| Model | Configuration | Accuracy | Precision | Recall | F1-Score |
| --- | --- | --- | --- | --- | --- |
| ResNet18 | 2D (F0 only) | 0.80 | 0.81 | 0.80 | 0.80 |
| ResNet50 | 2D (F0 only) | 0.82 | 0.82 | 0.82 | 0.81 |
| 3D-ResNet18 | 3D (7 planes) | **0.87** | **0.87** | **0.87** | **0.87** |
| 3D-ResNet50 | 3D (7 planes) | 0.86 | 0.86 | 0.86 | 0.86 |

**Fig. 5.**
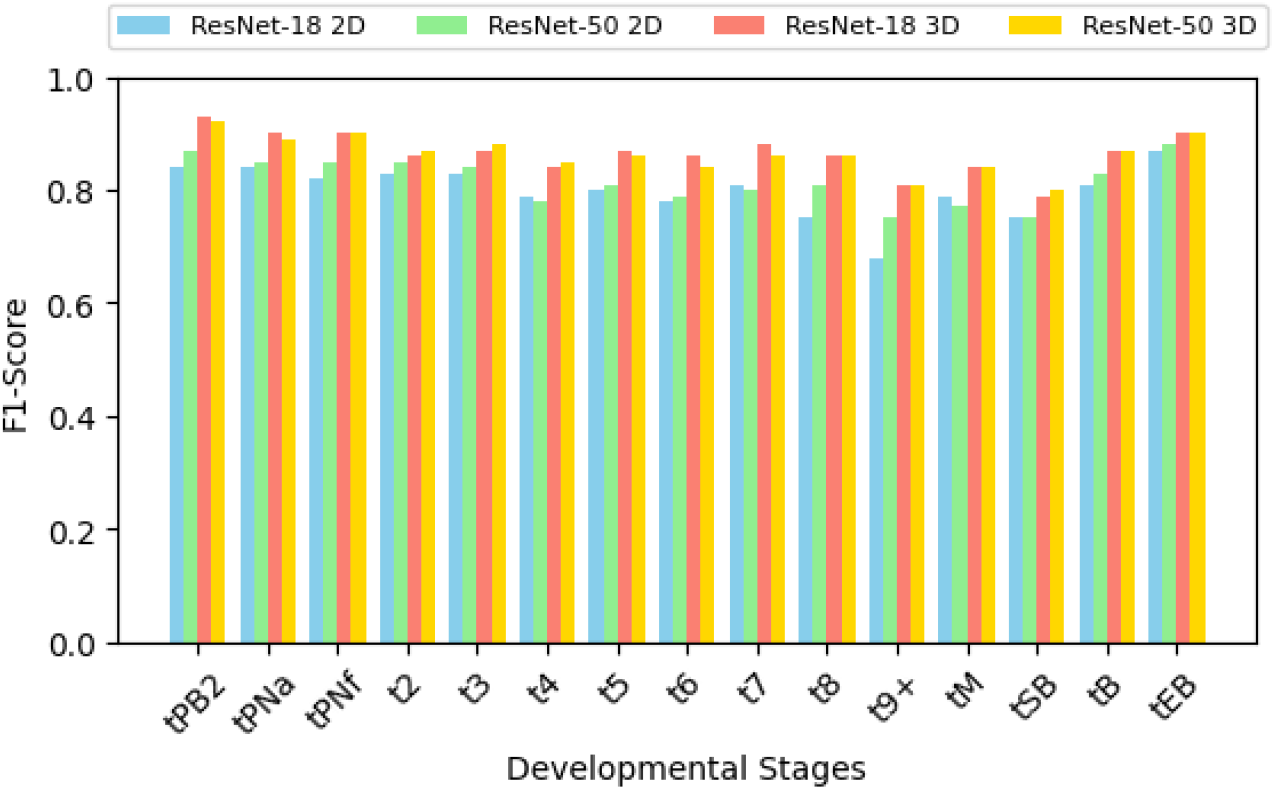
Illustration of F1-Score Across Developmental Stages for 2D and 3D Models.

### 4.5 Explainability

After applying the GradCAM XAI method described in 3.3, we perform a visual assessment of the saliency maps, to compare the models’ performances in capturing the most important regions for each prediction. We note that for the 3D-input cases, the saliency “volumes” produced by the models are in fact one 2D map, duplicated for each focal plane. While this specificity is being analyzed and corrected accordingly, in the present study we select only one slice of the saliency volumes and display it alongside the F0 input image, for comparison with 2D-models results.

On a global scale, we find satisfying maps across all models, and no obvious training issue is reported. For ResNet18 models, only one region is displayed, highlighting the cells, as expected in our understanding of the classification task (Fig. 6). On ResNet50 maps, we often find that the cells border is considered more relevant to the networks (Fig. 6 (b)), a characteristic that remains viable for prediction. These cases are ideal, but some mistakes can also be observed when navigating between models and classes.

**Fig. 6.**
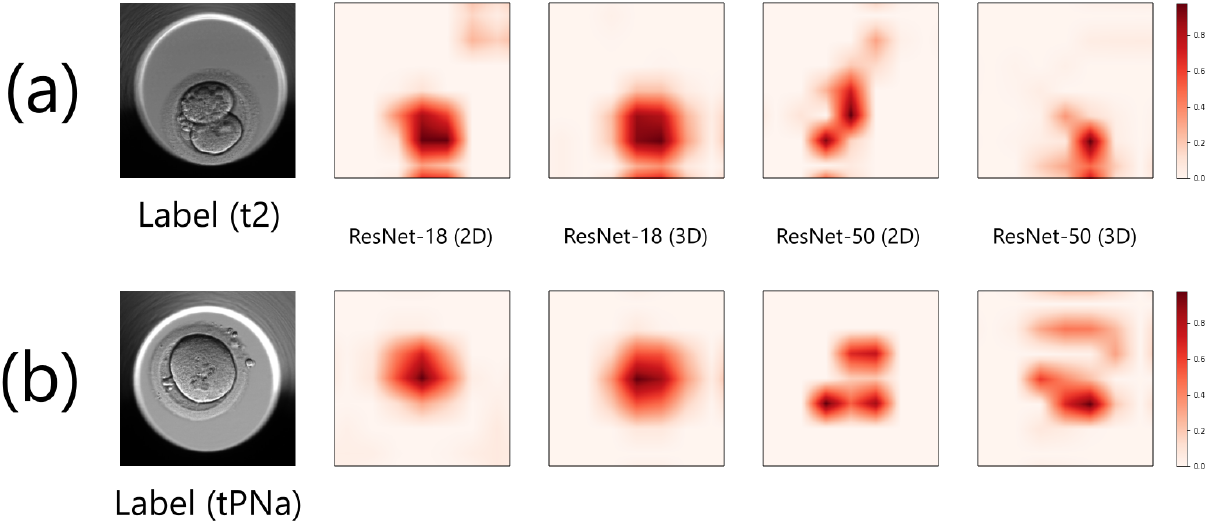
Examples for most satisfying GradCAM saliency maps, on all trained models. (a) All models can highlight the embryo cells, and (b) ResNet50 models often show the cells’ edges as most relevant.

#### ResNet18 models

In multiple saliency maps of both 2D and 3D models, in addition to the cells region, we can find the upper right corner of the image being strongly highlighted, as in Fig 7 (a). This area, outside the embryo itself, is clearly irrelevant for the growth classification, and its presumed relevance is most probably caused by a bias during training. However, this issue remains minor, since it rarely obfuscates the cells area from being also highlighted. Even when these failed explanations happen, and the cells are missed by one ResNet-18 model (Fig. 7 (b),(c)), the other architecturally similar network (2D or 3D) is able to display the correct relevant region in its stead, showing interesting complementary abilities.

**Fig. 7.**
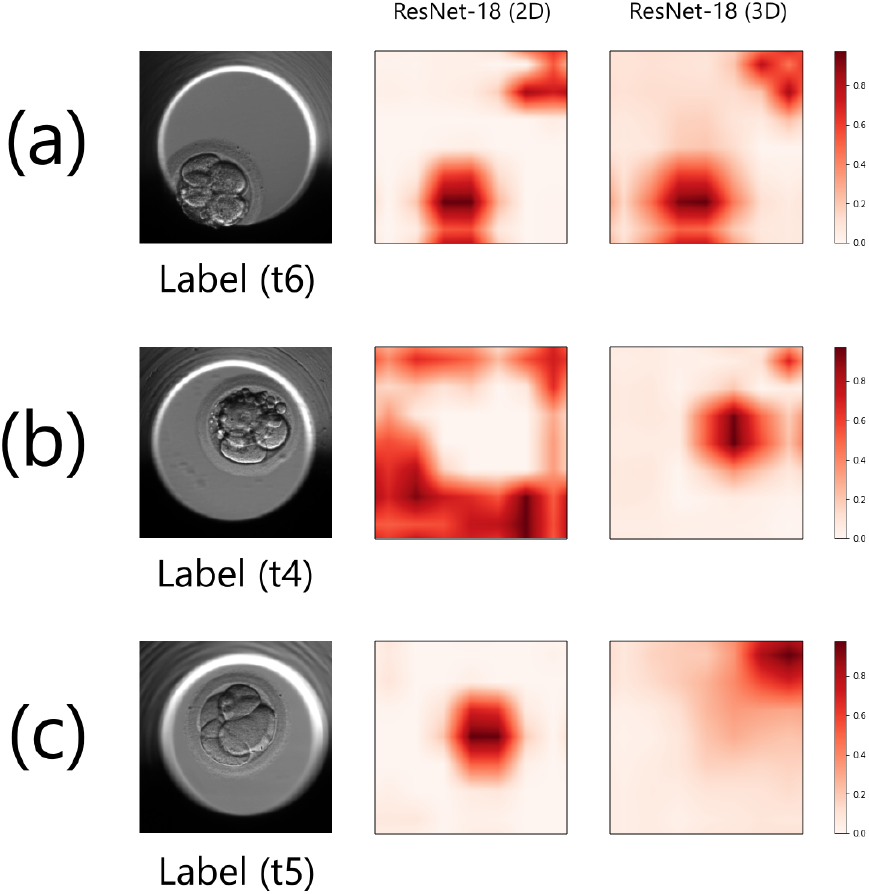
Mistakes examples in GradCAM saliency maps, for ResNet18 models: (a) the image upper right corner may also be highlighted; (b) and (c) when the cells are missed by one model, the other still performs well.

#### ResNet50 models

Unlike the smaller ResNet18 versions, ResNet50 models appear to be generally unfocused when explaining their predictions. In Fig. 8 (a) and (b), examples of saliency maps show most of the input image considered as relevant, including the border of the embryo’s cells. This observation can also lead to a complete miss (Fig. 8 (c)), when the entirety of the cells envelope is ignored, whereas the rest of the image is highlighted. In these situations, we often find both models being faulty at the same time, demonstrating a weaker learning of features importance than ResNet18 based models.

**Fig. 8.**
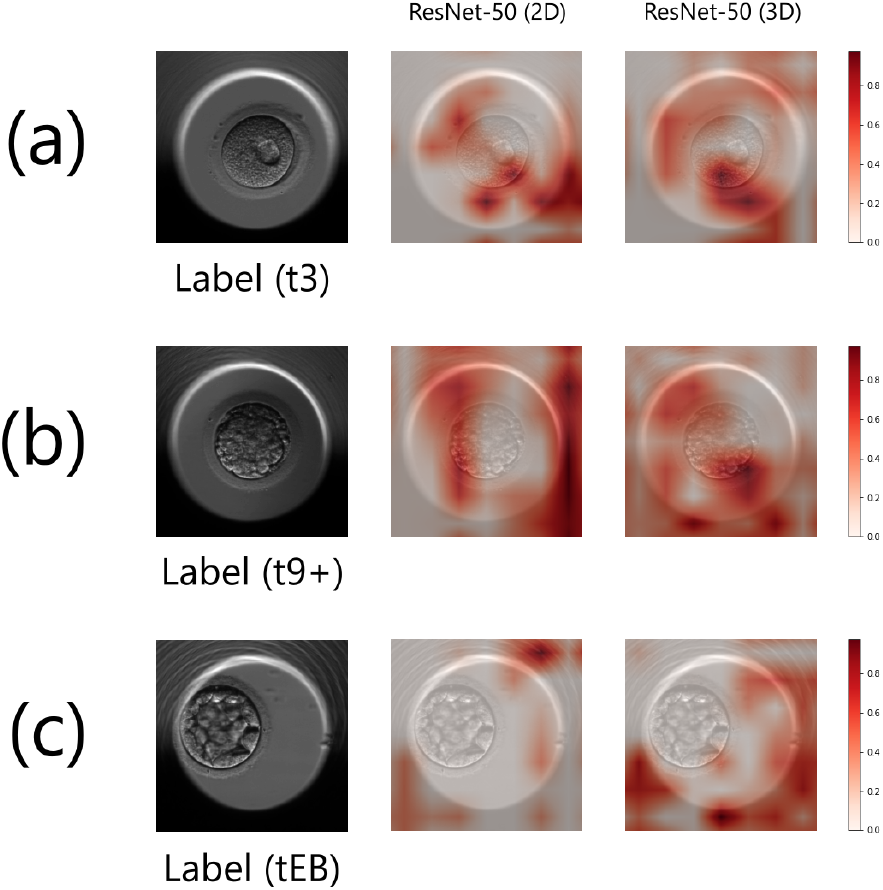
Mistakes examples in GradCAM saliency maps, for ResNet50 models: (a) and (b) despite highlighting the cells border, saliency maps are often noisy; (c) complete misses may happen for both models at the same time. For visualization purposes, the maps have been overlaid on the input image.

### 4.6 Discussion

Based on our expectations of relevant input areas, and the quality variance across saliency maps, ResNet18 models seem to have the best understanding of how to classify the embryos during their growth (ResNet50 models are too frequently unfocused). Moreover, there is no clear explainability improvement when using multiple focal planes, as both models have already satisfying reasoning abilities. When combined with the metrics performances in Table 4.4, we conclude that the 3D version of ResNet18 is the most promising model tested yet. A combination with its 2D counterpart may be beneficial for explainability purposes, but will prove costly in terms of inference time.

Regarding our attempt to explain the models reasoning using saliency concepts, the provided conclusions are in a preliminary state. In addition to the improvements proposed in 4.5, these results can be completed by integrating other XAI methods [17], with different features like pixel-wise attributions. For comparing the trained models, the use of quantitative metrics is also recommended [18], as visual assessment remains subjective.

## 5 Conclusion and perspectives

This study demonstrates that incorporating multi-focal plane imaging significantly improves embryo stage classification performance using deep learning models. The 3D ResNet architectures outperformed their 2D counterparts, particularly in complex developmental stages where depth information is crucial. Grad-CAM analysis confirmed that the models, especially ResNet18 variants, focused on relevant morphological features, though further refinement of explainability remains needed.

These results support the value of volumetric modeling in embryo assessment and open perspectives for integrating full 3D time-lapse sequences to capture both spatial and temporal dynamics of development. This combined approach could further enhance the model’s ability to detect subtle morphological changes over time.

Finally, data collection and expansion of multi-focal time-lapse datasets are ongoing. Future work will focus on validating the models on larger and clinically diverse cohorts, improving explainability with quantitative methods, and exploring computational optimizations for clinical deployment.

## Declarations

- **Funding:** No funding was received for this work.
- **Conflict of interest/Competing interests:** The authors declare no conflicts of interest.
- **Ethics approval and consent to participate:** Not applicable.
- **Consent for publication:** All authors consent to the publication of this work.
- **Data availability:** The dataset is accessible via https://doi.org/10.5281/zenodo.6390798
- **Author contributions:**
  - A. BS was responsible for coding, conducting experiments, and drafting an initial version of the manuscript.
  - W. M contributed to the explainability analysis and the interpretation of results.
  - S. H-C assisted with coding the approaches.
  - JM. G reviewed and approved the final manuscript.
  - M. EB contributed to the interpretation of the results and reviewed the manuscript.
  - F. M reviewed and approved the final manuscript.
  - A. P reviewed and approved the final manuscript.
  - MEA. B and N. S led the analysis, contributed to the reflection on the choice of approaches, directed the overall work, wrote the manuscript, and reviewed the paper.

